# FURNA: a database for function annotations of RNA structures

**DOI:** 10.1101/2023.12.19.572314

**Authors:** Chengxin Zhang, P. Lydia Freddolino

## Abstract

Despite the increasing number of 3D RNA structures in the Protein Data Bank, the majority of experimental RNA structures lack thorough functional annotations. As the significance of the functional roles played by non-coding RNAs becomes increasingly apparent, comprehensive annotation of RNA function is becoming a pressing concern. In response to this need, we have developed FURNA (<u>Fu</u>nctions of <u>RNA</u>s), the first database for experimental RNA structures that aims to provide a comprehensive repository of high-quality functional annotations. These include Gene Ontology terms, Enzyme Commission numbers, ligand binding sites, RNA families, protein binding motifs, and cross-references to related databases. FURNA is available at https://seq2fun.dcmb.med.umich.edu/furna/ to enable quick discovery of RNA functions from their structures and sequences.

## Introduction

Advances in experimental RNA structure determination methods, particularly Cryo-EM [1], have resulted in over 1600 RNA chains being deposited into the Protein Data Bank (PDB) database [2]. Despite these strides, functional annotations of experimental RNA structures are glaringly absent in both the PDB and secondary databases. The PDB database merely includes the bare minimum annotations for RNA chains, such as their names, lengths, and species. Downstream databases like NAKB (formerly known as NDB) [3], DNATCO [4], and BGSU RNA [5] offer more annotations for base pairing, backbone torsions, and 3D motifs, yet the annotation of the RNAs’ biological roles is still wanting. The MeRNA database [6] was probably the only database dedicated to function (in this case, metal ion binding sites) for experimental RNA structures, but it has long been defunct and was limited to 256 RNAs and their binding to metal ions. Simultaneously, recent studies have confirmed that many non-coding RNAs play vital roles in numerous biological events, particularly those involved in gene expression regulations [7], making RNA structures ideal targets for drug design [8]. This fresh understanding underscores the importance of annotating RNA functions for the RNA biology community.

In contrast to the stark lack of a functional database for RNA structures, several databases to annotate protein functions have already been established. Databases such as PDBsum [9] and SIFTS [10] annotate protein chains in the PDB using Gene Ontology (GO) terms and Enzyme Commission (EC) numbers by mapping PDB chains to UniProt [11] proteins and InterPro [12] families. The PDBbind-CN [13], BindingDB [14], and Binding MOAD [15] databases collect protein-ligand interactions with known affinity data. The PDBe-KB [16] database features ligand binding sites and post-translational modification sites for all PDB proteins. The FireDB [17] and IBIS [18] database curate protein-ligand interaction data in the PDB. Most recently, BioLiP2 [19] was developed as a comprehensive database covering almost all functional aspects of PDB proteins, including GO terms, EC numbers, ligand binding sites, binding affinities, and cross-reference to external databases.

Inspired by BioLiP2, we created FURNA, the first database in the field to offer comprehensive functional annotations for all RNA chains in the PDB database. Function annotations in FURNA include GO terms, EC numbers, Rfam [20] RNA families, RNA motifs for protein binding, species, literature, and cross references to external databases like PDBsum, NAKB, DNATCO, BGSU RNA, ChEMBL [21], DrugBank [22], ZINC [23], and RNAcentral [24]. Unlike protein-ligand interaction databases such as BioLiP, FireDB, and PDBbind-CN, which consider receptor-ligand contacts within each asymmetric unit, FURNA determines RNA-ligand interactions based on the biological assembly (i.e., biounit). This approach situates RNA-ligand interactions within the context of its quaternary structure. FURNA is available both as an open-source software package and as a browsable and searchable web service at https://seq2fun.dcmb.med.umich.edu/furna/.

## Results

### Overall statistics

At the time of writing this manuscript (Oct 2023), FURNA includes 16154 RNAs involved in 380680 ligand-RNA interactions; the online version of the database is updated on a weekly basis. Among these interactions, 186025, 138245, 31659, 24056 and 695 are interactions with metal ions, proteins, “regular” small molecule compounds excluding metal ions, other RNAs, and DNAs, respectively. Unlike BioLiP, FURNA does not attempt to exclude “biologically irrelevant” RNA-associated molecules from the database apart from removal of water molecules. This is because the biological relevance of ligands, especially metal ions, are less clearly defined for RNAs than for proteins. For example, calcium ions (Ca^2+^) are usually biologically irrelevant artifacts added for purification and/or crystallization purposes for proteins, but they are used to substitute magnesium ions (Mg^2+^) that are critical to maintain the folding of RNAs in pre-catalytic states [25]. Similarly, while potassium ions (K^+^) are a simple buffer additive for many proteins, they critical for the folding of many large RNAs where potassium ions stabilize juxtaposition of nucleotides with large sequence separation by neutralizing charge density [26]. This is why a significant portion (48.9%) of ligand-RNA interactions in FURNA are metal ions, among which 91.4% are magnesium ions, which are the most critical ion for RNA folding (**Fig 1A**). For non-ion small molecule ligands, the most frequent compound is osmium (III) hexammine (**Table S1**), which is a crystallization additive used to determine the RNA structure by multiwavelength anomalous diffraction (MAD) phasing [27].

**Fig 1.**
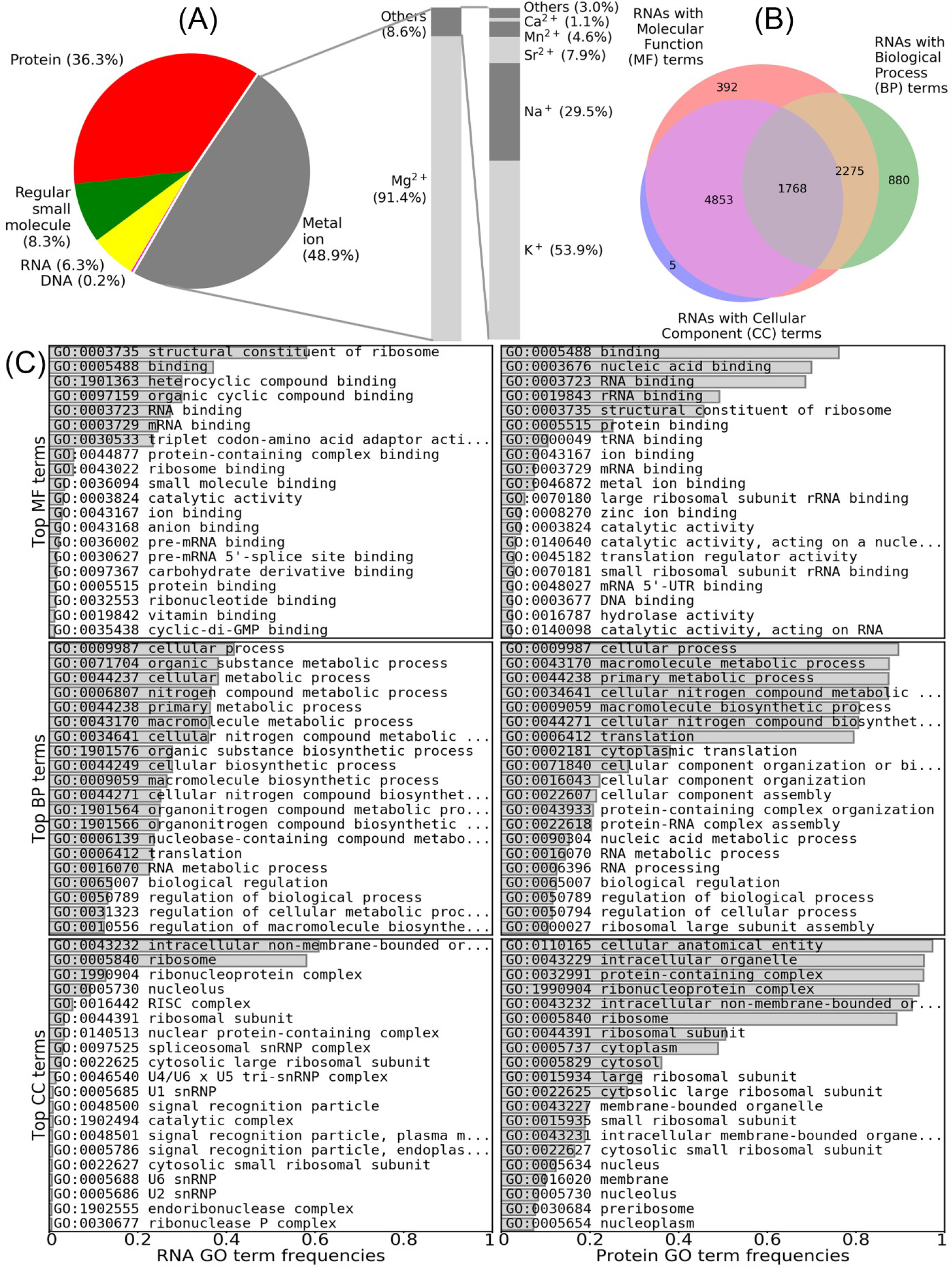
Overall statistics for RNAs and ligand-RNA interactions in FURNA. (A) Pie chart for the breakdown of ligand types in ligand-RNA interactions. (B) Venn graph showing the numbers of RNAs with GO terms in the MF, BP and/or CC aspects. (C) Top GO terms in RNAs and proteins collected by FURNA.

Among the 10561 RNAs with GO annotations in FURNA, 9288, 5311 and 7014 have annotations in Molecular Function (MF), Biological Process (BP), and Cellular Component (CC) aspects, respectively (**Fig 1B**). Out of the RNAs with MF terms, 58.0% are rRNAs (denoted by GO:0003735 “structural constituent of ribosome”) and 23.5% are tRNAs (indicated by GO:0030533 “triplet codon-amino acid adaptor activity”) (**Fig 1C**). This suggests that the distribution of RNA families among experimentally determined RNA structures is highly biased, consistent with MF annotations for RNA-binding proteins where GO:0019843 “rRNA binding” and GO:0000049 “tRNA binding” are among the most common GO terms. It is worth noting that, on average, the similarity of BP GO term annotations between an interacting RNA-protein pair is significantly higher than a random RNA-RNA pair or a random RNA-protein pair (**Figure S1**, Mann-Whitney U test p-value<1E-300 and p-value=1.0E-20, respectively). This suggests that RNA-protein interactions will be useful for RNA BP term prediction, similar to the utility of protein-protein interactions in protein function prediction [28-30].

### Web Interface

The FURNA website provides three primary interfaces: SEARCH, BROWSE, and DOWNLOAD. The functionalities of these interfaces are elaborated upon below.

#### BROWSE

Each entry in the FURNA database represents one RNA chain in the PDB. For each of these RNA chains, the BROWSE interface displays the PDB ID and chain ID, resolution, EC number, GO terms, RNAcentral accessions, Rfam families, species, PubMed citations, and protein-binding motifs found in the ATtrRACT [31] database (**Fig 2A**). Additionally, if the RNA chain has a non-water ligand, the BROWSE interface also provides information on the ligand ID, the chain and residue sequence number of the ligand, the ligand-binding nucleotides on the query RNA, and the biological assembly information where interaction with that ligand was retrieved.

**Fig 2.**
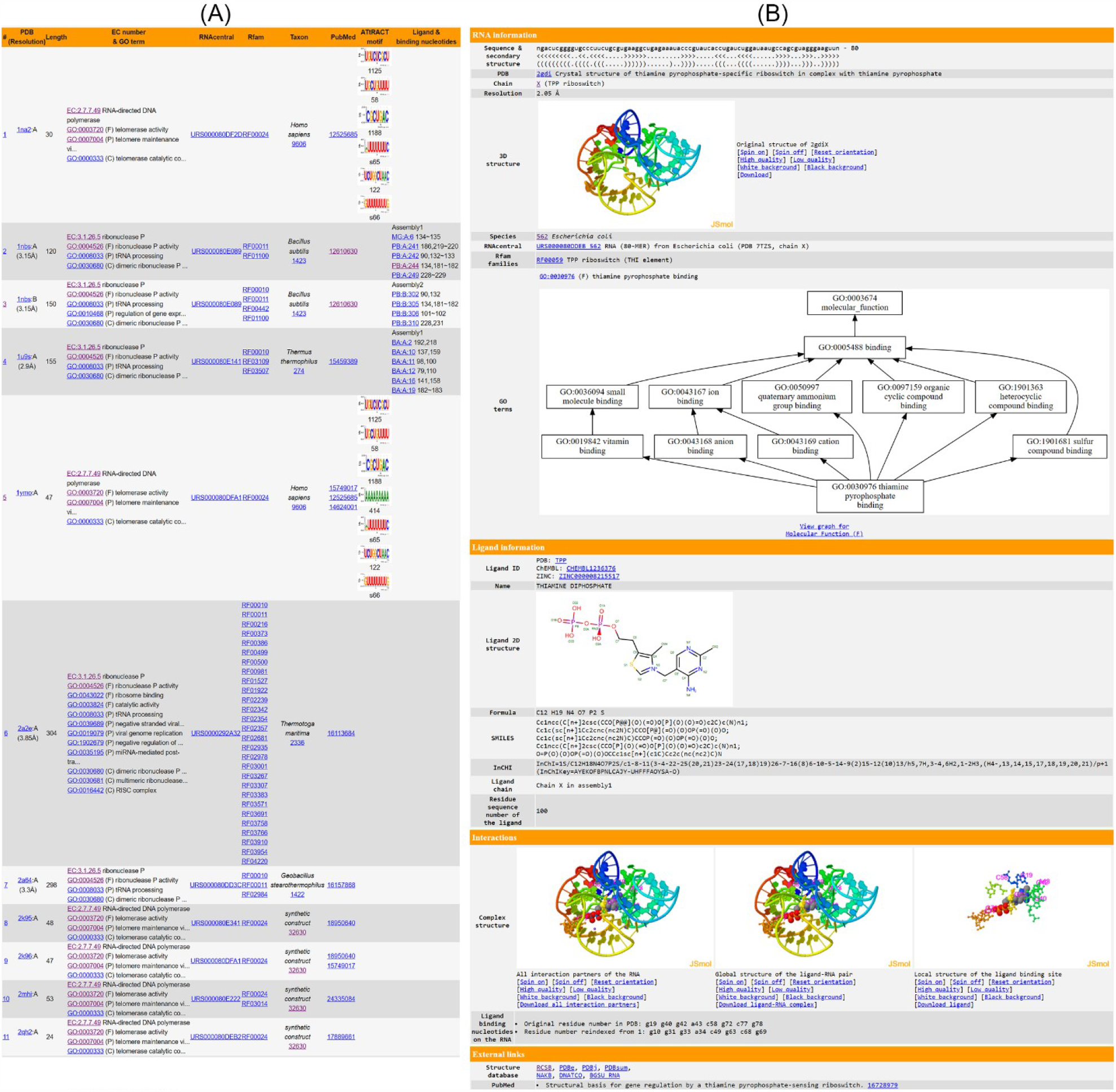
Browsing function annotations in FURNA. (A) Summary table. (B) Individual web page for small molecule-RNA interaction between thiamine diphosphate and TPP riboswitch (PDB 2gdi chain X, https://seq2fun.dcmb.med.umich.edu/furna/pdb.cgi?pdbid=2gdi&chain=X&lig3=TPP&ligCha=X&ligIdx=1).

Individual pages, accessed by clicking on the ligand in the last column of the BROWSE table, offer detailed structure and function information of each ligand-RNA interaction. These individual pages include the 3D structures of the RNA chain on its own, the full biological assembly, the RNA-ligand pair, and the local structure of the ligand binding site. These are displayed via four JSmol [32] applets. Where available, GO terms for Molecular Function, Biological Process, and Cellular Component are presented in three directed acyclic graphs created using Graphviz [33], illustrating the relationships among different GO terms. The GO terms, as well as EC numbers, are also listed as tables. Additional information is also provided, including the RNA sequence and secondary structure, resolution, the structure’s name, species, ATtRACT motifs, PubMed citations, and crosslinks to other databases. In case of a small molecule ligand or an ion, the page exhibits the 2D diagram, ligand IDs (including PDB CCD ID, ChEMBL ID, DrugBank ID, and ZINC ID), the chemical formula, ligand name, and linear descriptions of the molecules (**Fig 2B**).

For RNA, DNA, or protein ligands, additional details such as the sequence, name, and species, as well as relevant function annotations such as GO terms and EC numbers, are provided when available (**Figure S2**). For FURNA entries without ligand interactions, the structure and function details of the RNA can be viewed by clicking on the first column of the BROWSE interface.

#### SEARCH

The “SEARCH” interface provides four methods to explore FURNA: ‘Search by name’, ‘Quick sequence search (via BLAST)’, ‘Sensitive sequence search (via Infernal)’, and ‘Search by structure’. Firstly, users can query FURNA using PDB ID, PDB chain ID, ligand ID (as defined by the 3-letter code in the PDB database’s Chemical Compound Dictionary), ligand name, RNAcentral accession, Rfam family, EC number, GO term, ATtRACT motif, taxonomy, PubMed ID, or any combination of these. Secondly, FURNA can employ NCBI BLAST+ to search its entries using RNA, DNA, or protein sequences through a local non-redundant sequence database where identical sequences are merged into the same entry. In the search results, both representative hits found in the non-redundant database and members from the same sequence clusters are displayed (**Fig 3A**). Thirdly, to address the issue of a BLAST search’s low sensitivity for nucleic acid sequences, FURNA offers an alternative, more sensitive RNA sequence search option using Infernal (See Materials and Methods, **Fig 3B-C**). Lastly, users can search the tertiary structure of a query RNA (in PDB format) through the FURNA database using US-align (See Materials and Methods).

**Fig 3.**
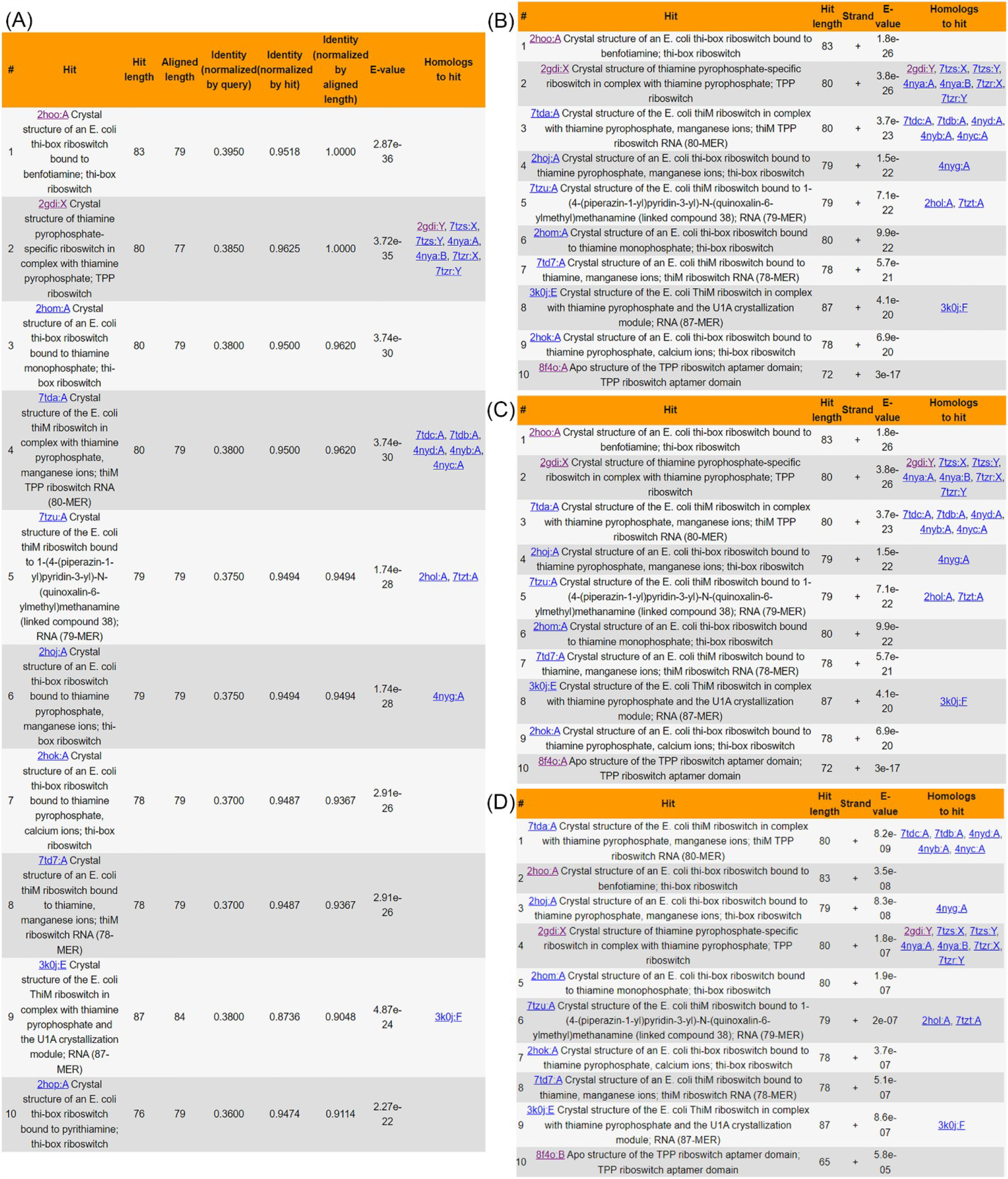
FURNA database sequence search results for TPP riboswitches in different bacteria. (A-B) Top 10 BLAST (A) and Infernal (B) search hits for the *E. coli* thiM riboswitch. (C-D) Top 10 Infernal search results for ykoF riboswitch (C) and tenA riboswitch (D) from *B. subtilis*. BLAST search result is not shown for the *B. subtilis* riboswitches because there are no BLAST hits.

#### DOWNLOAD

All data from FURNA can be downloaded in bulk through the “DOWNLOAD” interface. Functional annotations for each RNA chain and each ligand-RNA interaction are available in tab-separated tables. The FASTA sequences of RNAs, plus those of RNA-binding proteins and RNA-binding DNAs, are also provided. The coordinates of the RNAs and all non-water ligands are supplied in PDB format files. Furthermore, the link to the source codes for database curation and website display is also located on this page.

### Case study on TPP riboswitches

To illustrate FURNA’s utility in RNA function annotation, we conducted a case study involving the TPP (thiamin pyrophosphate) binding riboswitches, also known as the THI element or Thi-box riboswitch. This well-known family of riboswitches binds to thiamine pyrophosphate (TPP) to regulate the expression of its downstream gene [34, 35]. In *Escherichia coli*, one such riboswitch is located upstream of the Hydroxyethylthiazole kinase (*thiM*) coding sequence [36, 37] (**Fig. 2B, Table S2**). Unsurprisingly, both BLAST (**Fig. 3A**) and Infernal (**Fig. 3B**) searches of the *E. coli* TPP riboswitch through FURNA return hits for many TPP riboswitches, including those from *E. coli*. Similar results can be obtained by searching the region upstream of the *thiM* gene of *Siccibacter turicensis*, which also belongs to the Enterobacteriaceae family (**Table S2**).

Based on gene function and the general prevalence of Thi-box riboswitches, we suspected the presence of riboswitches at several locations in *Bacillus subtilis*, e.g., one situated upstream of the coding sequence of the HMP/thiamine-binding protein (*ykoF*) and the other situated upstream of the aminopyrimidine aminohydrolase (tenA, **Table S2**). Indeed, the *tenA* riboswitch has been previously reported [38], whereas a riboswitch upstream of *ykoF* has not, to our knowledge, been previously reported in the literature, although its presence is indicated in the RNAcentral database (https://rnacentral.org/rna/URS000005CA97). When using FURNA to perform a BLAST sequence search of the putative *B. subtilis* TPP riboswitches, no hits are returned, not to the *E. coli thiM* riboswitch. This outcome is not unexpected considering *E. coli* and *B. subtilis* are gram-negative and gram-positive bacteria, respectively, and have evolved separately for billions of years. In contrast, a sensitive Infernal search using either of the potential *B. subtilis* TPP riboswitches does yield hits to other TPP riboswitches, including the *E. coli thiM* riboswitch (**Fig. 3C-D**). These findings highlights FURNA’s capability for function annotations of low-homology RNAs using its sensitive sequence search option, providing a unified interface for obtaining functional information on a new RNA of interest.

## Discussion and Conclusions

We introduce FURNA, the first comprehensive structure database for ligand-RNA interactions and RNA function annotations. Compared to existing RNA structure and function databases, FURNA stands out in several ways. Firstly, it is the only database to utilize standard function vocabularies (GO terms and EC numbers) for the annotation of RNA tertiary structures. Secondly, it outlines ligand-RNA interactions based on biological assembly, which enhances the investigational context of interactions within the complete RNA-containing complex. Thirdly, FURNA offers user-friendly database search capabilities at varying levels of sensitivity, ensuring its relevance in annotating even remote RNA homologs. Fourthly, its data curation code is modular and fully open source, thereby simplifying regular data updates and future development. These unique aspects of FURNA position it as a valuable resource for the biological community, aiding in summarizing known RNA biological functions, creating functional hypotheses for poorly characterized RNAs, and developing new algorithms for ligand-RNA docking, virtual screening, and structure-based RNA function annotation. Nonetheless, FURNA does present a challenge in its lack of a clear definition of the biological relevance of ligand-RNA interactions, an issue we plan to address in our future work.

## Materials and Methods

Each entry in FURNA corresponds to one RNA chain in the PDB database. To this end, we first download the mmCIF files of all structures containing nucleic acid from the PDB database and split them into individual chains using a modified version of the BeEM tool [39]. An RNA chain is defined by possessing more ribonucleotides than deoxyribonucleotides and amino acids. RNA chains with ten or more nucleotides become entries in FURNA, but oligo-ribonucleotide fragments with fewer than ten nucleotides are only included as “ligands” if they bind to an RNA chain with ten or more nucleotides. The curation of an RNA chain involves several steps: annotating GO terms and EC numbers, mapping RNA-protein binding motifs, extracting RNA-ligand interactions, and assigning RNA secondary structures.

### GO term and EC number annotation

We employ two complementary strategies to obtain GO terms for an RNA chain. First, we search the RNA sequence against the most current version of the Rfam database (Rfam 14.9, with covariance models for 4108 families) using Infernal [40], utilizing the parameters: `cmsearch --cpu 4 -Z 1 --toponly`. Then, we transfer the GO terms related to each Rfam family hit (http://current.geneontology.org/ontology/external2go/rfam2go) to the query RNA. Second, we map RNA chains to RNAcentral sequences based on the mapping file provided by RNAcentral (http://ftp.ebi.ac.uk/pub/databases/RNAcentral/current_release/id_mapping/database_mappings/pdb.tsv). If the RNAcentral entry has GO terms in the Gene Ontology Annotation (GOA) project (http://ftp.ebi.ac.uk/pub/databases/GO/goa/UNIPROT/goa_uniprot_all.gaf.gz), we also transfer these GO terms to the FURNA entry. We utilize Graphviz [33] to plot the direct acyclic graphs showcasing the relationships among an RNA’s GO terms (including their parent terms). For the subset of RNAs with annotated catalytic activities, we convert their Enzyme Commission (EC) numbers from GO terms using the EC2GO mapping (https://www.ebi.ac.uk/GOA/EC2GO). For RNA-binding proteins, their UniProt accessions, GO terms and EC numbers are directly obtained through the SIFTS [10] database.

### RNA-protein binding motif mapping

To identify RNA motifs corresponding to known recognition sites for RNA-binding proteins, we download the position weight matrices (PWMs) for all 1583 protein-binding motifs from the latest ATtRACT database (version 0.99β). These motifs and the query RNAs collected by FURNA are grouped by species. Here, we extract the species information of an RNA chain from the respective mmCIF file, specifically from records such as “gene_src_ncbi_taxonomy_id”, “ncbi_taxonomy_id”, “pdbx_gene_src_ncbi_taxonomy_id”, or “pdbx_ncbi_taxonomy_id”. For any species that has at least one ATtRACT motif and one FURNA RNA chain, we download its transcriptome from the NCBI FTP site (ftp://ftp.ncbi.nlm.nih.gov/genomes/all/annotation_releases/) to determine its background distribution of the four nucleotide types (A, C, G, U). This background information is ascertained using the fasta-get-markov program of the MEME suite [41]. Subsequently, this background file is used by the FIMO program [42] of the MEME suite when it searches the motif PWMs against the FURNA RNAs with the parameters: `--norc --bfile`, to enable the motifs to align with the RNAs.

### Ligand-RNA interaction extraction

For each query RNA included in the FURNA database, we gather its interaction partners from the mmCIF format biological assembly file (i.e., biounit) that contains the pertinent RNA chain. As an example, the asymmetric unit of PDB 1a9n (the spliceosomal U2B”-U2A’ protein complex bound to a fragment of U2 small nuclear RNA) contains six chains, which comprises four protein chains (Chains A, B, C, and D) and two RNA chains (Chains Q and R). This PDB correlates with two different biological assemblies: assembly 1 includes chains A, B, and Q; assembly 2 incorporates chains C, D, and R. Consequently, to extract ligand-RNA interactions for 1a9n chain R, we only consider assembly 2.

Starting from the biological assembly file selected for a query RNA, we employ a modified version of the BeEM program [39] to split it into different chains. For each chain, we further split the macromolecule part and the small molecule parts, where the former and latter are labeled by numerical values and a period (“.”), respectively, in the “label_seq_id” record of the mmCIF file. Next, we collect all non-water ligands from all chains in the mmCIF file, including small molecules and metal ions, proteins, DNAs, and RNAs (excluding the query RNA). For each query RNA-ligand pair, we calculate all inter-molecular atomic contacts, i.e., atom pairs within the sum of the van der Waals radii plus 0.5 Å, among non-hydrogen atoms. We label a nucleotide on the query RNA as a ligand binding residue if it has two or more inter-molecular atomic contacts with a ligand. We group any collection of two or more ligand binding residues for the same query RNA-ligand pair into a binding site. Ligands without a binding site are excluded.

For a small molecule ligand, we extract the name, synonyms, chemical formula, and linear descriptions (including SMILES, InChI, and InChIKey) from the Chemical Component Dictionary (CCD) provided by the PDB database. We perform mappings from PDB ligand IDs (i.e., CCD IDs) to ligand IDs in the ChEMBL, DrugBank, and ZINC databases using the UniChem database [43]. For protein ligands, we retrieve their GO terms, EC numbers, species, and UniProt accessions from the SIFTS [10] database. For DNA ligands, we retrieve the species from the mmCIF file of the asymmetric unit, similar to how we obtain species information for RNA chains.

### RNA secondary structure assignment

FURNA assigns RNA secondary structures in dot-bracket format for canonical base pairs (Watson-Crick pairs and G:U Wobble pairs) in the experimental 3D structure, using two complementary methods: CSSR [44] and DSSR [45]. CSSR is our in-house program, optimized for coarse-grained RNA structures. It can assign secondary structures even when the nucleotides have missing atoms. Conversely, DSSR only functions when the nucleobase of the nucleotide is fully atomic and its RMSD to the standard nucleobase conformation is less than 0.28 Å. Due to these stringent requirements, DSSR-assigned secondary structures might have missing positions compared to the input RNA. To ensure consistency between DSSR input and output, we utilize Arena [46] to fill in missing atoms and rectify unphysical nucleobase conformations for all RNA chains before we execute the DSSR assignment. For an RNA-RNA interaction involving two RNA chains, we assign secondary structures to both the individual RNAs and the RNA pair.

### Infernal database construction

For users to perform sensitive Infernal searches of query RNA sequences through FURNA, a database in the Infernal [40] format must be pre-constructed. To accomplish this, we first obtain a non-redundant set of RNAs, which is generated by collapsing multiple FURNA RNAs with identical sequences into one entry. For each RNA in the non-redundant set, the CSSR-assigned secondary structure in dot bracket is collected, and any pseudoknots present in the secondary structures are removed. Subsequently, the secondary structure and sequence are converted by the “cmbuild” tool of the Infernal package into the uncalibrated Infernal format covariance model. This covariance model is then calibrated by the “cmcalibrate” tool of the Infernal package. The calibrated covariance models for all non-redundant FURNA RNA chains are concatenated into the Infernal format database. This database can be utilized by the “cmscan” tool of the Infernal package, allowing a user to perform Infernal searches of query RNA sequences through FURNA.

### US-align database construction

Since conducting a tertiary structure search of all RNA chains in FURNA is more time-consuming than a sequence search, two procedures are implemented to reduce the size of the structure database used for US-align search. First, the non-redundant set of RNAs with non-identical sequences is isolated, from which the coordinates of the C3’ atoms are extracted. The exclusion of atoms other than C3’ does not affect US-align, which only considers C3’ atoms for RNA structure alignment. Second, we utilize the qTMclust tool [47] from the US-align package to cluster the structures of the non-redundant RNAs. This results in a set of representative RNA structures with a pairwise TM-score [48] less than 0.5. These representative RNA structures form the US-align database. When a user carries out an RNA structure query through the FURNA website, this query structure will be searched using US-align through the database of representative structures to report the top 100 hits with the highest TM-scores. Meanwhile, the RNAs belonging to the same structure clusters will also be reported.

## Acknowledgements

We thank Dr Xiaoqiong Wei for insightful discussions. This work was directly supported by NIH R01 AI134678 (to PLF). This work also used the Advanced Cyberinfrastructure Coordination Ecosystem: Services & Support (ACCESS) program, which is supported by National Science Foundation (Grant nos. 2138259, 2138286, 2138307, 2137603, and 2138296).

## Author contributions

C.Z. conceived the project and developed the method. C.Z. and P.L.F designed the experiment, performed the data analysis, and wrote the manuscript.

